# *Chrna5* is essential for a rapid and protected response to optogenetic release of endogenous acetylcholine in prefrontal cortex

**DOI:** 10.1101/2020.05.10.087569

**Authors:** Sridevi Venkatesan, Evelyn K. Lambe

## Abstract

Optimal attention performance requires cholinergic modulation of corticothalamic neurons in the prefrontal cortex. These pyramidal cells express specialized nicotinic acetylcholine receptors containing the α5 subunit encoded by *Chrna5*. Disruption of this gene impairs attention, but the advantage α5 confers for the detection of *endogenous* cholinergic signaling is unknown. To ascertain this underlying mechanism, we used optogenetics to stimulate cholinergic afferents in prefrontal cortex brain slices from compound-transgenic wild-type and *Chrna5* knockout mice of both sexes. These electrophysiological experiments identify that *Chrna5* is critical for the *rapid onset* of the postsynaptic cholinergic response. Loss of α5 slows cholinergic excitation and delays its peak, and these effects are observed in two different optogenetic mouse lines. Disruption of *Chrna5* does not otherwise perturb the magnitude of the response, which remains strongly mediated by nicotinic receptors and tightly controlled by autoinhibition via muscarinic M2 receptors. However, when conditions are altered to promote *sustained* cholinergic receptor stimulation, it becomes evident that α5 also works to protect nicotinic responses against *desensitization*. Rescuing *Chrna5* disruption thus presents the double challenge of improving the onset of cholinergic signaling without triggering desensitization. Here, we identify that an agonist for the unorthodox α-α nicotinic binding site can allosterically enhance this cholinergic pathway considered vital for attention. Minimal NS9283 treatment restores the rapid onset of the postsynaptic cholinergic response without triggering desensitization. Taken together, this work demonstrates the advantages of speed and resilience that *Chrna5* confers on endogenous cholinergic signaling, defining a critical window of interest for cue detection and attentional processing.

**Significance statement:** The α5 nicotinic receptor subunit (*Chrna5*) is important for attention, but its advantage in detecting endogenous cholinergic signals is unknown. Here, we show that α5 subunits permit rapid cholinergic responses in prefrontal cortex and protect these responses from desensitization. Our findings clarify why *Chrna5* is required for optimal attentional performance under demanding conditions. To treat the deficit arising from *Chrna5* disruption without triggering desensitization, we enhanced nicotinic receptor affinity using NS9283 stimulation at the unorthodox α-α nicotinic binding site. This approach successfully restored the rapid-onset kinetics of endogenous cholinergic neurotransmission. In summary, we reveal a previously unknown role of *Chrna5* as well as an effective approach to compensate for genetic disruption and permit fast cholinergic excitation of prefrontal attention circuits.

## Introduction

The medial prefrontal cortex (PFC) is essential for working memory and top-down attention (Goldman-Rakic, 1995; Miller and Cohen, 2001; Dalley et al., 2004). Cholinergic neuromodulation of the prefrontal cortex by projections from the basal forebrain is required for attention (Dalley et al., 2004). These projections release acetylcholine during successful cue detection and performance of sustained attention tasks (Himmelheber et al., 2000; Parikh et al., 2007; Gritton et al., 2016; Howe et al., 2017). Corticothalamic neurons in layer 6 are excited by such acetylcholine release (Kassam et al. 2008; Hedrick and Waters, 2015; Sparks et al., 2018). They express the specialized α5 nicotinic receptor subunit encoded by *Chrna5* (Wada et al., 1990; Winzer-Serhan and Leslie, 2005) in addition to the more commonly-expressed α4 and β2 subunits of high-affinity nicotinic receptors. The α5 subunit has been shown to increase calcium permeability and sensitivity of α4β2* nicotinic receptors to acetylcholine in cell systems (Tapia et al., 2007; Kuryatov et al., 2008) and to boost the response to exogenous stimulation *ex vivo* in mouse prefrontal cortex (Bailey et al., 2010; Tian et al., 2011). However, the impact of the α5 subunit on synaptic cholinergic neurotransmission in prefrontal cortex is unknown.

Behavioural and genetic evidence suggest that the α5 nicotinic subunit plays a role in attention and more generally in prefrontal executive function. Mice lacking *Chrna5* display attention deficits in the 5 choice serial reaction time test, exhibiting a failure to detect cues when the task difficulty is increased to make cue duration shorter (Bailey et al., 2010). Work in rats has also confirmed that *Chrna5* is important for performing demanding attention tasks (Howe et al., 2018). Human polymorphisms in *Chrna5* that affect receptor functionality are associated with a cognitive phenotype that increases early experimentation with smoking and risk for addiction (Bierut et al., 2008), as well as increased risks of schizophrenia, cognitive impairments and attention deficit hyperactivity disorder (Hong et al., 2011; Schuch et al., 2016; Han et al., 2019).

Despite the link with attention and prefrontal cognitive processes, the role of *Chrna5* in responding to endogenous acetylcholine release has yet to be examined. Most α5 characterization relies on heterologous expression systems and not the natural environment of nicotinic receptors in neurons (Baenziger and daCosta, 2013). While there is tight control over trafficking of cholinergic receptors (Matta et al., 2017), the lack of validated antibodies for immuno-electron microscopy means the relationship between α5 and cholinergic afferents is unknown. Functional examination lacks a specific pharmacological tool to discriminate α5 subunit-containing nicotinic receptors. Studies characterizing cholinergic functionality in α5 knockout mice have used exogenous applications of acetylcholine that differ vastly from the rapid timescales of endogenous neurotransmission (Parikh et al., 2007). Thus, there is a gap in our understanding of the role of *Chrna5* in cholinergic modulation of attention circuits. The development of optogenetic tools to specifically express channelrhodopsin in cholinergic neurons (Zhao et al., 2011a; Hedrick et al., 2016) allows us to overcome this gap and measure the function of the α5 subunit in endogenous cholinergic modulation.

To probe the advantage α5 confers on endogenous cholinergic signaling, we optogenetically stimulated cholinergic afferents in prefrontal brain slices of compound transgenic wild-type and α5 knockout mice. Concurrent whole cell electrophysiology shows that the α5 subunit is essential for achieving rapid kinetics of cholinergic neurotransmission. Under conditions of prolonged stimulation, the α5 subunit preserves the synaptic cholinergic response from desensitization. A pharmacological approach targeting the recently-discovered α-α acetylcholine binding site on nicotinic receptors (Harpsøe et al., 2011; Wang et al., 2015; Mazzaferro et al., 2017) rescues onset-kinetics of the cholinergic response after α5 disruption. Recent perspectives on cholinergic modulation have sought to shed light on the temporal scales of cholinergic signaling (Disney and Higley, 2020; Sarter and Lustig, 2020). In this context, our work reveals a critical and specialized role for the α5 nicotinic receptor subunit in initiating rapid cholinergic signaling.

## Methods

### Animals

In order to elicit endogenous acetylcholine release optogenetically and examine responses in α5 wild-type (α5WT) and α5 knockout (α5KO) littermate mice, we created two compound transgenic mouse lines. Mouse crosses are illustrated in the respective figures using these mice. The first was achieved by crossing ChAT-ChR2 (JAX: 014546) (Zhao et al., 2011b) with α5KO mice (Salas et al., 2003) to achieve parents. These mice were crossed with α5HET mice to generate α5WT and α5KO ChAT-ChR2/+ mice. We independently verified the results of optogenetic cholinergic stimulation using a different line of mice to express channelrhodopsin in cholinergic neurons. ChAT-IRES-Cre (JAX: 031661) and Ai32 mice (JAX: 012569) were each crossed with α5HET mice and their offspring were crossed with each other to generate littermate α5WT and α5KO ChAT-IRES-Cre/+ Ai32/+ mice. All animals were bred on a C57BL/6 background. Both male and female animals age >P60 were used. Mice were weaned at P21, separated based on sex, and group housed (2-4 mice per cage) and given ad libitum access to food and water on a 12-h light/dark cycle with lights on at 7 AM. Guidelines of the Canadian Council on Animal Care were followed, and all experimental procedures were approved by the Faculty of Medicine Animal Care Committee at the University of Toronto. A total of 92 mice were used for the entire study, with similar numbers of males and females.

### Electrophysiology

Animals were anesthetized with an intraperitoneal injection of chloral hydrate (400 mg/kg) and then decapitated. The brain was rapidly extracted in ice cold sucrose ACSF (254 mM sucrose, 10 mM D-glucose, 26 mM NaHCO_3_, 2 mM CaCl_2_, 2 mM MgSO_4_, 3 mM KCl and 1.25 mM NaH_2_PO_4_). Coronal slices 400 μm thick of prefrontal cortex (Bregma 2.2 - 1.1) were obtained on a Dosaka linear slicer (SciMedia, Costa Mesa, CA, USA). Slices were allowed to recover for at least 2 hours in oxygenated (95% O_2_, 5% CO_2_) ACSF (128 mM NaCl, 10 mM D-glucose, 26 Mm NaHCO_3_, 2 mM CaCl_2_, 2 mM MgSO_4_, 3 Mm KCl, and 1.25 mM NaH_2_PO_4_) at 30°C before being used for electrophysiology.

For whole cell patch clamp electrophysiology, brain slices were transferred to a chamber mounted on the stage of a BX51WI microscope (Olympus, Tokyo, Japan) and constantly perfused with oxygenated ACSF at 30°C. Layer 6 pyramidal neurons were patched based on their morphology and the proximity to white matter. The recording electrodes (2 - 4 MΩ) were filled with patch solution composed of 120 mM potassium gluconate, 5 mM KCl, 10 mM HEPES, 2 mM MgCl_2_, 4 mM K_2_-ATP, 0.4 mM Na_2_-GTP and 10 mM sodium phosphocreatine, pH adjusted to 7.3 using KOH. Data were acquired with Multiclamp 700B amplifier at 20 kHz with Digidata 1440A and pClamp 10.7 acquisition software (Molecular devices). All recordings were compensated for the liquid junction potential (14 mV). Layer 6 pyramidal responses were examined in voltage-clamp at -75 mV and in current-clamp at rest or starting from -70 mV.

There are two distinct population of pyramidal neurons in layer 6 which differ in their spiking pattern to current injection – regular spiking corticothalamic neurons and doublet spiking corticocortical neurons (Kumar and Ohana, 2008; Ledergerber and Larkum, 2010; Thomson, 2010). As previously reported using endogenous (Hedrick and Waters, 2015) and exogenous acetylcholine (Yang et al., 2019), we found that the corticocortical neurons exhibit a purely muscarinic receptor mediated hyperpolarization, while regular spiking corticothalamic neurons exhibit nicotinic receptor mediated depolarization. Therefore, our experiments focused on the regular spiking corticothalamic pyramidal neurons to characterize the role of α5 subunit containing nicotinic receptors in the response to endogenous acetylcholine release.

### Optogenetics

To excite channelrhodopsin containing axonal fibers, blue light (473 nm) was delivered in brief pulses (5 ms) with an LED (Thorlabs, 2 mW) through the 60X objective lens. 8 pulses of light, 5 ms each were delivered in a frequency accommodating manner, starting at 50 Hz and ending in 10 Hz to stimulate the cholinergic axons (Fig 1A – experimental schematic). This paradigm was intended to replicate the accommodating firing pattern of cholinergic neurons (Unal et al., 2012). As indicated, a subset of experiments alternatively used only a single pulse of light (5 ms).

**Figure 1:**
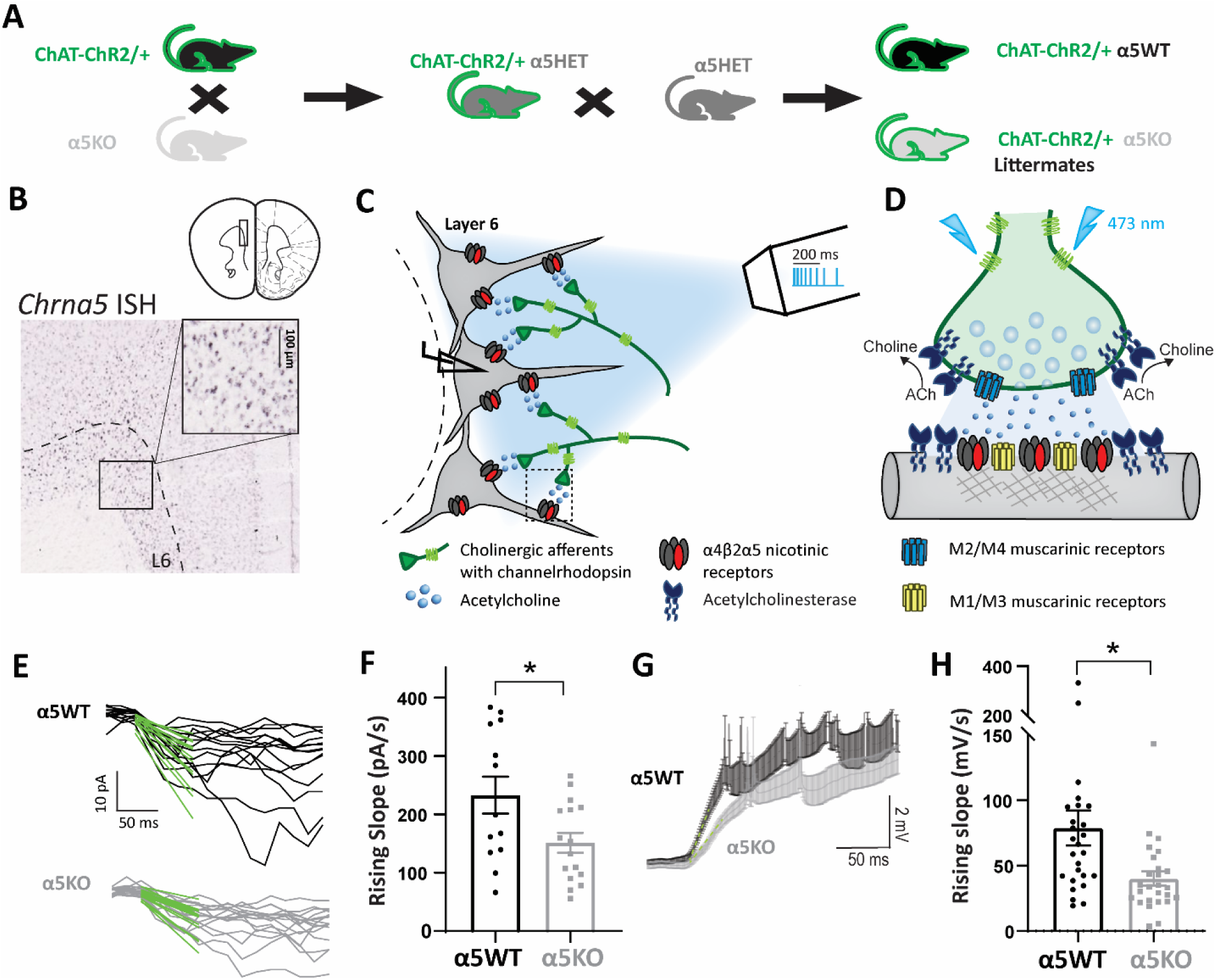
*Chrna5* is critical for maintaining rapid onset of the response to optogenetic acetylcholine release. **A**, Schematic showing mouse crosses to obtain littermate wildtype (α5WT) and α5 knockout (α5KO) ChAT-ChR2 mice. **B**, *In situ* hybridization for *Chrna5* mRNA in mouse prefrontal cortex shows expression in layer 6 neurons (Image from Allen Institute). Schematic showing coronal slice of mouse brain is adapted from (Paxinos and Franklin, 2004). **C**, Schematic depicting experimental approach of whole cell patch clamping of layer 6 pyramidal neurons in brain slices to measure responses to optogenetic release of acetylcholine from cholinergic afferents expressing channelrhodopsin. Optogenetic stimulation pattern of 8 pulses (5 ms) in frequency accommodating manner (50 - 10 Hz) is shown with scale bar (legend and detailed illustration of cholinergic synapses in D). **D**, Schematic showing optogenetic stimulation of a cholinergic synapse causing acetylcholine release onto layer 6 pyramidal neurons in the prefrontal cortex. Different effectors shown are postsynaptic nicotinic receptors with the *Chrna5* subunit ((α4)_2_(β2)_2_α5 receptors), M1/M3 muscarinic receptors, presynaptic M2/M4 autoinhibitory muscarinic receptors and acetylcholinesterase. **E**, Fast rising phase of cholinergic responses in voltage-clamp (−75 mV) in WT (top) and α5KO (bottom) and linear fit (green) to the first 50 ms of the response from light onset. **F**, Bar graph showing the rising slope (pA/s) of the current determined from the linear fit in WT and α5KO (unpaired t-test: *t*_(27)_ = 2.34, \**p* = 0.02). α5KO show slower onset of cholinergic responses. **G**, Average current-clamp response of WT and α5KO (*n* = 26 cells each) layer 6 pyramidal neurons shows a slower rise in α5KO. **H**, Bar graph showing the rising slope (mV/s) of the depolarization determined from the linear fit in WT and α5KO (unpaired t-test: *t*_(50)_ = 2.68, \*\**p* = 0.001). α5KO show slower onset of cholinergic responses.

### Pharmacology

Atropine (200 nM, Sigma) was applied to block muscarinic receptors. AFDX-116 (300 nM, Tocris) was used to block M2 muscarinic receptors. Dihydro-β-erythroidine (DhβE, 3 μM, Tocris) was used to block β2 containing nicotinic receptors. CNQX (20 μM, Alomone), APV (50 μM, Alomone) and picrotoxin (50 μM, Alomone) were used to block glutamate and GABA-A receptors. Diisopropylfluorophosphate (DFP, 20 μM, Toronto Research Chemicals) was used to block acetylcholinesterase and induce spillover of acetylcholine. Nicotine (100 nM, Sigma) was used for desensitization experiments. All experiments with nicotine and DFP were done in the presence of atropine to isolate the nicotinic response. The selective agonist of the α-α binding site, NS9283 (100nM, Tocris) was used to restore rapid rise time of the nicotinic response in layer 6 neurons of the α5KO mice.

### Analysis and statistics

Analysis of cholinergic responses was performed in Clampfit 10.2 (Molecular Devices) and Axograph. Raw traces were used for calculating the rising slope of the voltage-clamp response within 50 ms of light onset to get accurate measurement of the fast onset kinetics. Downsampled traces were used to fit double exponentials to the cholinergic responses and for representation. Exponential and linear fits to the responses were performed on Axograph. Magnitude of the cholinergic responses in voltage-clamp were determined by the peak current (pA) as well as the charge transferred (pC) into the cell which is measured as the area under the current response for 1 s starting from the light onset. Graphpad Prism 8 was used for statistical analysis and plotting graphs. Genotype differences in parameters of the cholinergic responses between α5WT and α5KO were compared with two-tailed unpaired *t*-tests where applicable. Effect of pharmacological manipulations in WT and α5KO were compared with Sidak’s post-hoc test, and repeated measures/ paired *t*-tests were used if the recordings were obtained from the same cell pre and post pharmacology. When comparing the effect of a drug (e.g. nicotine) across time between α5WT and α5KO, data were analyzed using 2-way repeated measures ANOVA (or mixed-effects analysis in the case of missing time points for some cells) and Sidak’s multiple comparisons test used to compare α5WT and α5KO at each time point or between the baseline condition and different time points of drug application within each genotype. All ANOVAs were performed with the Geisser Greenhouse correction for sphericity. *P* values < 0.05 were considered statistically significant. Data are reported as mean ± SEM.

## Results

### Chrna5 is critical for maintaining a rapid-onset response to endogenous acetylcholine

In order to assess the contribution of the α5 nicotinic subunit to endogenous cholinergic neurotransmission, we bred compound transgenic mice to achieve channelrhodopsin-labelled cholinergic fibers in littermate α5WT and α5KO mice, as illustrated in **Figure 1A**.We recorded regular-spiking pyramidal neurons in layer 6 to obtain a population of neurons known to have nicotinic acetylcholine receptors enriched for α5 (Bailey et al., 2010), as illustrated by *Chrna5* expression in Figure 1B. Figure 1D shows a schematic of the hypothesized components of the cholinergic synapse onto these layer 6 pyramidal neurons based on available data (Levey et al., 1991; Zhang et al., 2002; Hedrick and Waters, 2015; Sparks et al., 2018). Since basal forebrain cholinergic neurons innervating the prefrontal cortex are thought to fire in brief bursts with spike frequency accommodation (Unal et al. 2012; Lee et al. 2005), we chose a pattern of optogenetic stimulation to mimic burst firing with 8 pulses of blue light (473 nm) delivered in a frequency-accommodating manner as illustrated in the schematic in Fig 1C.

Our examination of layer 6 neuron cholinergic responses to optogenetic stimulation revealed that there were distinct differences in the onset kinetics between the α5WT and α5KO. We fit a line to the fast-rising phase of the cholinergic responses (initial 50 ms) to calculate the rising slope of the response (Fig 1E & G). The rising slope of the cholinergic current is significantly smaller in α5KO neurons (151 ± 17 pA/s) compared to the WT (233 ± 32 pA/s; unpaired t-test: *t*_(27)_ = 2.34, \**p* = 0.02; *N* = 7 mice per genotype, Fig 1F). This difference in rising kinetics was also reflected in the current-clamp responses, with the α5KO having a significantly slower rising slope (40 ± 5 mV/s), ∼ 50% when compared to the WT neurons (79 ± 14 mV/s; unpaired t-test: *t*_(50)_=2.68, \*\**p* = 0.001; Fig 1H). This slower onset of cholinergic responses in the α5KO translates to a significant delay in cholinergic activation induced spiking (delay in onset of first spike in α5KO: 87 ± 39 ms; unpaired t-test comparing onset of first spike in WT and α5KO: *t*_(14)_ = 2.25, \**p* = 0.04). For both voltage-clamp and current-clamp examination, the peak response amplitudes themselves were not significantly different between the two genotypes (Peak current: unpaired t-test: *t*_(26)_ = 0.38, *p* = 0.70; Peak depolarization: unpaired t-test: *t*_(35)_ = 1, *p* = 0.34). There were no sex differences nor sex-by-genotype interactions on any measure of the endogenous cholinergic response (data not shown). Furthermore, genotype differences in response onset kinetics were observed in the absence of genotype differences in passive electrophysiological properties **(Table 1).** These results indicate that the α5 subunit is critical for rapid onset of cholinergic activation in layer 6 of the prefrontal cortex.

**Table 1:** Electrophysiological properties of α5WT and α5KO layer 6 pyramidal neurons. The intrinsic properties of layer 6 pyramidal neurons from Figure 1 did not differ statistically between α5WT and α5KO.

| | $\alpha 5$WT<br>(7 mice) | $\alpha 5$KO<br>(7 mice) | unpaired t-test | Summary |
| --- | --- | --- | --- | --- |
| Resting membrane potential (mV) | $-87 \pm 1$ | $-87 \pm 1$ | $t_{(76)} = 0.1, p = 0.9$ | ns |
| Spike amplitude (mV) | $78 \pm 2$ | $76 \pm 2$ | $t_{(76)} = 0.5, p = 0.6$ | ns |
| Input resistance (MOhm) | $140 \pm 7$ | $153 \pm 12$ | $t_{(76)} = 0.9, p = 0.3$ | ns |
| Capacitance (pF) | $73 \pm 4$ | $65 \pm 2$ | $t_{(76)} = 1.6, p = 0.1$ | ns |
| Membrane time constant (ms) | $10 \pm 1$ | $10 \pm 1$ | $t_{(76)} = 0.5, p = 0.6$ | ns |

### Autoinhibitory control of layer 6 cholinergic release is strong and Chrna5-independent

Cholinergic synaptic transmission is likely to include activation of pre-and postsynaptic muscarinic receptors as shown in **Figure 2A**, and it has been suggested previously that α5 knockouts can show compensatory increases in muscarinic excitability (Tian et al., 2011). Therefore, we examined the contribution of muscarinic receptors in the response to endogenous cholinergic neurotransmission. Of note, the pan-muscarinic receptor antagonist atropine increased the magnitude of the cholinergic responses in both genotypes (2 way repeated measures ANOVA, Effect of atropine: *F*_(1,14)_ = 30.87, **** *p* < 0.0001, *N* = 5, 4 mice for α5WT, α5KO; Fig 2B & C). We hypothesized this overall response potentiation is due to the block of presynaptic autoinhibitory M2/M4 muscarinic receptors on cholinergic terminals. We tested this hypothesis by specifically blocking the M2 muscarinic receptor that is the main autoinhibitory receptor in the cortex (Levey et al., 1991; Zhang et al., 2002) using AFDX-116. Blocking M2 muscarinic receptors significantly potentiates the cholinergic response in both WT and α5KO (2 way repeated measures ANOVA: Effect of AFDX-116: *F*_(1,6)_ = 16.32, \*\**p* = 0.007) with no significant interaction between the effect of AFDX-116 and genotype (AFDX-116 x Genotype interaction : *F*_(1,6)_ = 3.50, *p* = 0.1, effect of genotype: *F*_(1,6)_ = 0.2, *p* = 0.7). These results suggest that cholinergic modulation of prefrontal layer 6 is under active regulation by presynaptic M2 muscarinic receptors in both genotypes.

**Figure 2:**
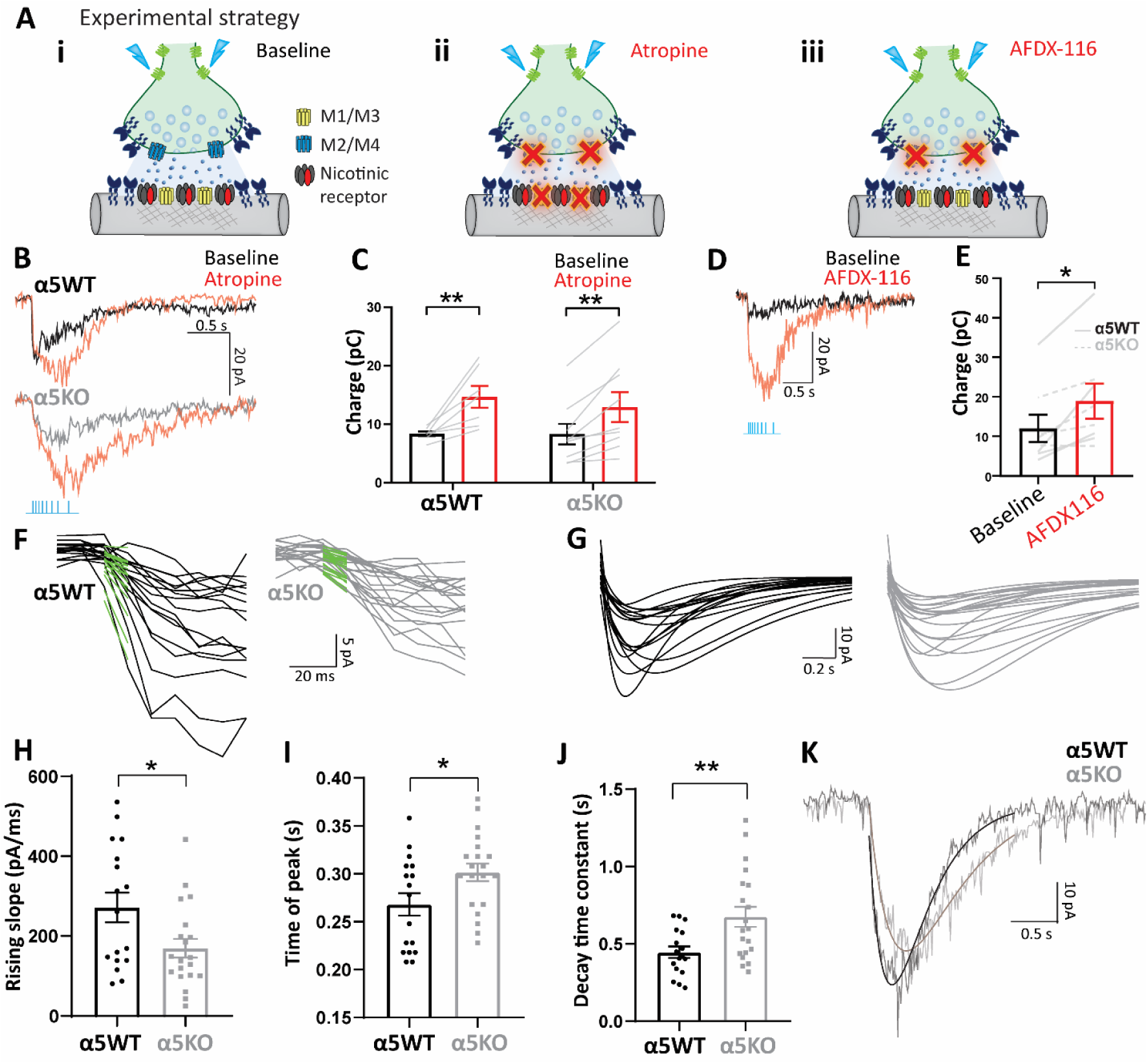
Autoinhibition of optogenetically-released acetylcholine by M2 muscarinic receptors is strong and *Chrna5*-independent. **A i-iii**, Experimental schematic illustrating cholinergic synaptic transmission at (i) baseline, (ii) when all muscarinic receptors both postsynaptic M1/M3 and presynaptic M2/M4 are blocked by atropine and (iii) when presynaptic autoinhibitory M2/M4 muscarinic receptors are selectively blocked by AFDX-116. **B**, Cholinergic response of a WT (top) and α5KO neuron (bottom) in voltage-clamp before and after the application of atropine (200 nM). **C**, Bar graph showing cholinergic current charge at baseline and after atropine in WT and α5KO. Atropine significantly increases cholinergic current charge (2-way RM ANOVA: Effect of atropine: *F*_(1,14)_ = 30.87, **** *p* < 0.0001) in both WT (Sidak’s post-hoc test: *t*_(14)_ = 4.3, \*\**p* = 0.002) and α5KO (Sidak’s post-hoc test: *t*_(14)_ = 3.5, \*\**p* = 0.006) to the same extent. **D**, Cholinergic response of a WT neuron in voltage-clamp before and after the application of M2 antagonist AFDX-116 (300 nM). **E**, Bar graph showing cholinergic current charge before and after the application of AFDX-116 in WT and α5 KO. Blocking presynaptic autoinhibitory muscarinic receptors is sufficient to significantly increase cholinergic response magnitude in both WT and α5KO (paired *t*-test: *t*_(7)_ = 3.47, \**p* = 0.01). **F**, Fast rising phase of cholinergic responses in WT (top) and α5KO (bottom) and linear fit (green) to the first 50 ms of the response from light onset after the application of atropine. **G**, Double exponential fits to cholinergic responses in WT (left) and α5KO (right) neurons. **H-J**, Bar graph showing the (H) rising slope (pA/s) of the current determined from the linear fit (unpaired t-test: *t*_(35)_ = 2.40, \**p* = 0.02) (I) Time of peak current (unpaired t-test: *t*_(35)_ = 2.27, \**p* = 0.03) and (J) Decay time constant determined from double exponential fits (unpaired t-test: *t*_(35)_ = 2.93, \*\**p* = 0.006) of the cholinergic responses in WT and α5KO. **K**, Example cholinergic response with exponential fits of a WT and α5KO neuron illustrates slower onset, delayed peak and slower decay in α5KO.

The genotype differences in the kinetics of the cholinergic responses were still evident following block of muscarinic receptors, with the α5KO neurons showing significantly slower rising slope (170 ± 23 pA/s) compared to the WT (272 ± 37 pA/s, unpaired t-test: *t*_(35)_ = 2.40, \**p* = 0.02; Fig 2F & H). The time of peak current, as measured from the exponential fits to the responses (Fig 2G & I) is significantly delayed by 33.5 ± 14.7 ms in the α5KO compared to the WT (unpaired t-test: *t*_(35)_ = 2.27, \**p* = 0.03; Fig 2I). In addition to this slower onset observed both at baseline and in the presence of atropine, we also find that the α5KO showed a significantly greater decay time constant (675 ± 65 ms) compared to WT neurons (446 ± 37 ms; unpaired t-test: *t*_(35)_ = 2.93, \*\**p* = 0.006; Fig 2J). Although the response kinetics were different between genotypes, the charge transfer did not differ by genotype either before or with atropine (Fig 2C) nor were there differences in the response pharmacology. Nicotinic receptor-mediated responses to acetylcholine release were completely eliminated in both WT and α5KO (98% reduction) by the β2 nicotinic receptor antagonist DhβE (2 way repeated measures ANOVA: Effect of DhβE : *F*_(1,4)_ = 38.96, \*\**p* = 0.003), with no significant interaction nor main effect of genotype (effect of genotype: *F*_(1,4)_ = 0.27, *p* = 0.6; genotype x DhβE interaction: *F*_(1,4)_ = 0.11, *p* = 0.8). This indicates that in the absence of the α5 subunit, the nicotinic receptors in the α5KO remain β2-containing receptors.

### Replication in a different optogenetic model: Chrna5 is required for rapid cholinergic kinetics

To test the robustness of the finding that α5KO mice show a selective deficit in kinetics of cholinergic activation, we repeated our experiments in a different line of mice targeting channelrhodopsin to cholinergic neurons. To create WT and α5KO offspring for these experiments, we bred compound crosses of α5het/ChAT-IRES-Cre and α5het/Ai32 mice as illustrated in **Figure 3**. It has been recently reported that this fate-mapping approach will include a subset of neurons which are only cholinergic transiently during development and release glutamate in the adult (Nasirova et al., 2019). Therefore, we examined all recordings obtained in response to optogenetic stimulation for evidence of fast glutamatergic EPSCs time locked to the stimulus onset and included only recordings without such light-evoked EPSCs. We additionally performed a subset of experiments in the presence of glutamate receptor blockers CNQX and APV and found no significant differences from the data acquired without glutamate blockers (2 way repeated measures ANOVA: effect of CNQX+APV: *F*_(1,7)_ = 2.49, *p* = 0.16).

**Figure 3:**
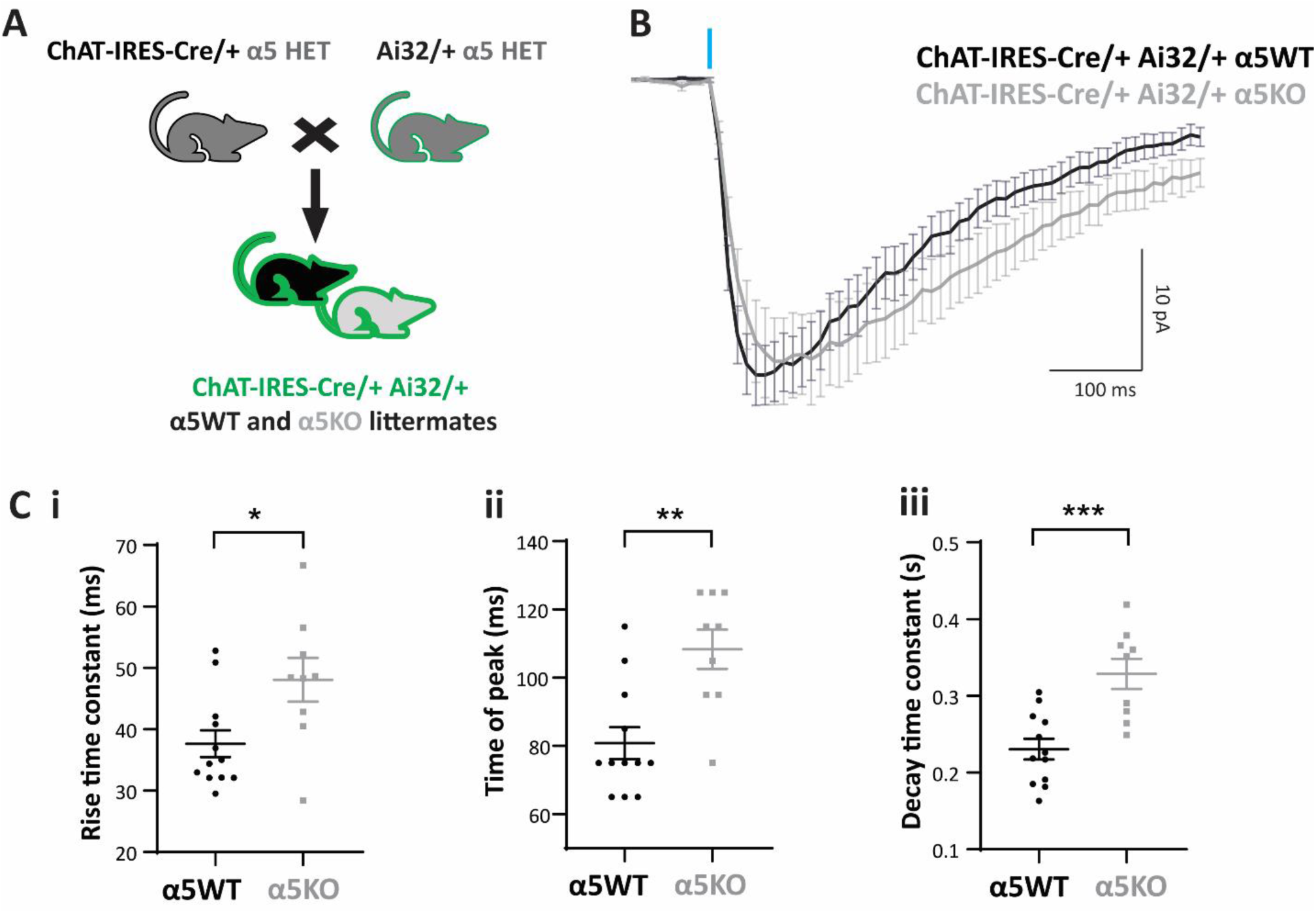
*Chrna5* maintains rapid cholinergic kinetics in a different optogenetic model. **A**, Schematic showing mouse crosses to obtain littermate wildtype (α5WT) and α5 knockout (α5KO) ChAT-IRES-Cre/+ Ai32/+ mice. **B**, Average cholinergic current obtained in response to one 5 ms flash of optogenetic stimulation in α5WT (*n* = 12) and α5KO (*n* = 9) neurons in the presence of atropine. **C i-iii**, Bar graph showing (i) rise time constant (unpaired t-test: *t*_(19)_ = 2.6, \**p* = 0.02) (ii) time of peak (unpaired t-test t_(19)_ = 3.74, \*\**p* = 0.001) and (iii) decay time constant (unpaired t-test: t_(29)_ = 3.26, \*\**p* = 0.003) in WT and α5KO. Cholinergic responses in the α5KO have a slower rise, delayed peak, and slower decay compared to WT.

The ROSA26 promoter in ChAT-IRES-Cre/+ Ai32/+ α5WT and α5KO expressed channelrhodopsin more strongly in the cholinergic afferents than the ChAT promoter, and a single pulse of light (5 ms) was sufficient to generate a cholinergic response of comparable magnitude to that examined in Fig 2. The average cholinergic current elicited in layer 6 pyramidal neurons by a single 5 ms pulse of light stimulation in α5WT (*n* = 12 cells) and α5KO (*n* = 9 cells) is shown in figure 3B (*N* = 4 mice per genotype). Consistent with results from the α5KO ChAT-ChR2 mice, the cholinergic response in the α5KO is delayed compared to WT, with the rise time constant significantly greater in α5KO (48 ± 11 ms) compared to WT (38 ± 8 ms; unpaired t-test: *t*_(19)_ = 2.6, \**p* = 0.02; Fig 3C i). Although the α5KO show slower rise, they attain a similar peak magnitude (28 ± 4 pA) as WT (29 ± 3 pA; unpaired t-test: *t*_(19)_ = 0.22, *p* = 0.8), but the α5KO peak occurs at a significantly delayed time point (delay in time of peak in α5KO: 27.5 ± 7 ms; unpaired t-test *t*_(19)_ = 3.74, \*\**p* = 0.001; Fig 3C ii). The α5KO also show significantly slower decay time constant compared to WT (α5WT: 353 ± 16 ms vs α5KO: 438 ± 20 ms, unpaired t-test: *t*_(29)_ = 3.26, \*\**p* = 0.003; Fig 3C iii). We are thus able to replicate the key deficits in cholinergic response kinetics observed in α5KO ChAT-ChR2 mice in α5KO ChAT-IRES-Cre/+ Ai32/+ mice. We conclude that *Chrna5* is essential to maintain the rapid onset of response to acetylcholine release in layer 6 pyramidal neurons.

### Chrna5 protects endogenous cholinergic signaling against desensitization

Acetylcholine levels in the prefrontal cortex increase during attention, arousal, exploration and other cognitive tasks, as well as during stress (Pepeu and Giovannini, 2004; Gritton et al., 2016; Teles-Grilo Ruivo et al., 2017a). To understand the consequences of prolonged acetylcholine presence occurring under situations of high cognitive effort, we blocked acetylcholine breakdown by inhibiting acetylcholinesterase irreversibly with diisopropylfluorophosphate (DFP). We measured layer 6 neuron responses to a train of optogenetic stimulation in WT and α5KO ChAT-ChR2 mice before and after acetylcholinesterase inhibition with DFP, in the continuous presence of atropine (**Figure 4A-C**). Blocking acetylcholinesterase causes the optogenetic cholinergic responses to nearly double in the WT (cholinergic current charge at baseline: 16 ± 2 pC, after DFP: 32 ± 5 pC; Sidak’s post hoc test: *t*_(58)_ = 2.92, \**p* = 0.01; *N* = 5 mice), but the increase was not significant in the α5KO (cholinergic current charge at baseline: 13 ± 2 pC, after DFP: 18 ± 3 pC; *t*_(58)_ = 0.85, *p* = 0.64; *N* = 3 mice; Fig 4C). Overall, there were significant main effects of DFP and genotype on the charge transfer from the optogenetic cholinergic response, (2-way ANOVA: effect of DFP: *F*_(1,58)_ = 7.29, \*\**p* = 0.009; effect of genotype: *F*_(1,58)_ = 5.90, \**p* = 0.02; DFP x genotype interaction: *F*_(1,58)_ = 2.34, *p* = 0.13). Post-hoc comparison shows that after acetylcholinesterase inhibition, the cholinergic charge transfer is significantly lower in α5KO (18 ± 3 pC) compared to WT (32 ± 5 pC, Sidak’s post hoc test: *t*_(58)_ = 3, \*\**p* = 0.008). These responses to endogenous acetylcholine release in the presence of cholinesterase inhibitors are reminiscent of genotype differences observed for direct responses to exogenous acetylcholine application (Bailey et al., 2010), where there is a prolonged high concentration of acetylcholine at the synapse due to saturation of acetylcholinesterase.

**Figure 4.**
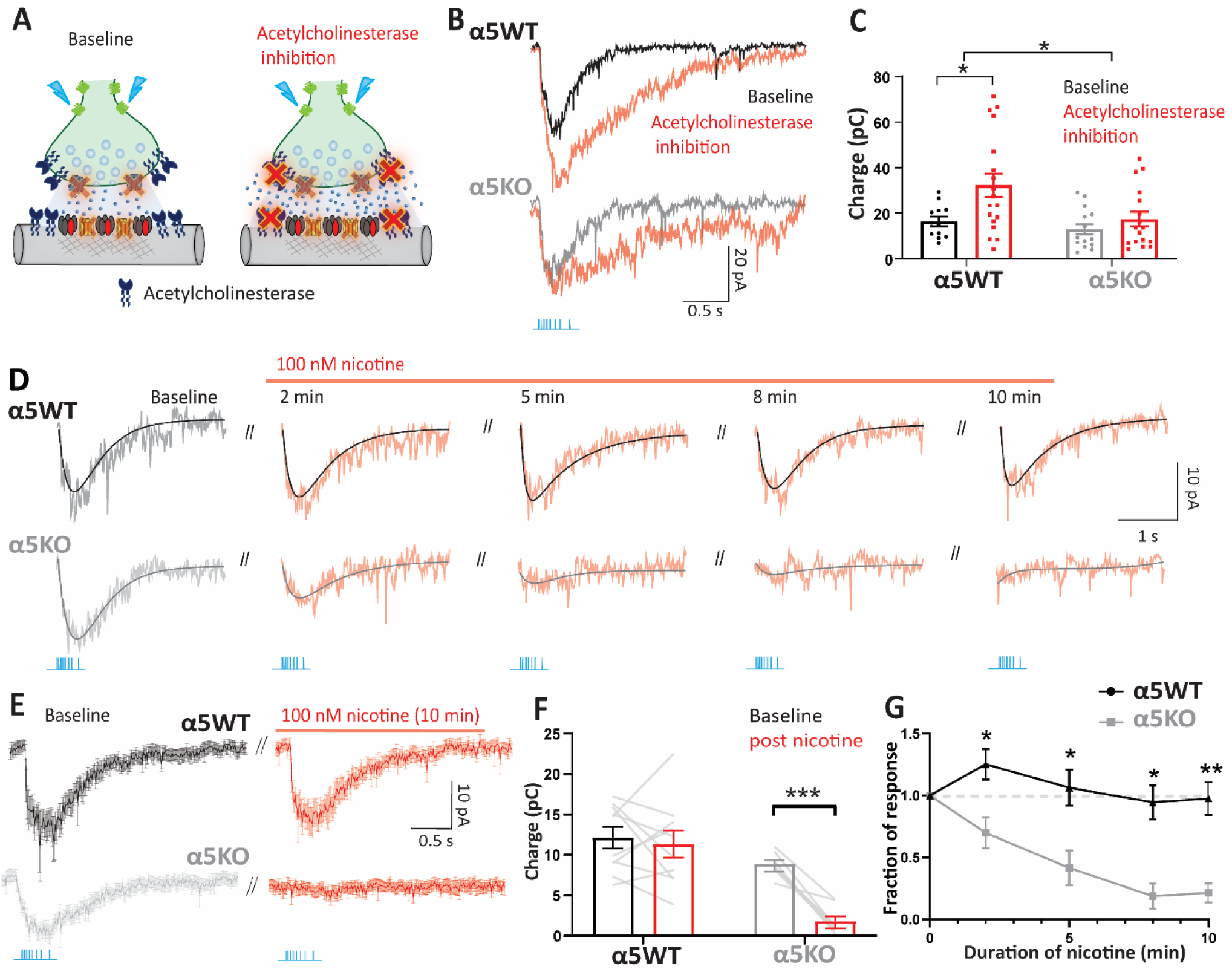
*Chrna5* protects endogenous cholinergic signaling against desensitization. **A**, Experimental schematic illustrating nicotinic synaptic transmission at baseline when muscarinic receptors are blocked by atropine and following the application of an acetylcholinesterase inhibitor (DFP) to prevent breakdown of acetylcholine. **B**, Nicotinic response of a WT (top) and α5KO neuron (bottom) in voltage-clamp before and after the application of the acetylcholinesterase inhibitor DFP (20 μM). **C**, Bar graph showing nicotinic current charge in WT and α5KO at baseline and in the presence of DFP. There was a significant effect of acetylcholinesterase inhibition on the cholinergic responses (2-way ANOVA: Effect of DFP: *F*_(1,58)_ = 7.29, \*\**p* = 0.009), but also a significant effect of genotype (*F*_(1,58)_ = 5.90, \**p* = 0.02). Cholinesterase inhibition caused a significant increase in the magnitude of cholinergic responses in WT (Sidak’s post hoc test: *t*_(58)_ = 2.92, \**p* = 0.01), but did not enhance cholinergic responses in the α5KO (*t*_(58)_ = 0.85, *p* = 0.64). **D**, Optogenetically evoked nicotinic responses with their exponential fits in α5WT and KO at different time points during the application of 100 nM nicotine for 10 minutes. **E**, Average nicotinic current in response to optogenetic acetylcholine release in α5WT (*n* = 5) and α5KO (*n* = 6) neurons before and after 10 min nicotine. **F**, Bar graph showing nicotinic current charge before and after 10 min nicotine in WT and α5KO (2-way RM ANOVA: Genotype x nicotine interaction: *F*_(1,11)_ = 12.56, \*\**p* < 0.01, Sidak’s post hoc comparison of baseline and post-nicotine responses in WT: *p* = 0.9, in α5KO: \*\*\**p* < 0.001). **G**, Time course of change in endogenous nicotinic response as nicotine is applied (Sidak’s posthoc test comparing WT and α5KO: \**p* < 0.05 \*\**p* < 0.01).

We hypothesized that this *Chrna5* genotype difference in the ability of the optogenetic response to withstand prolonged exposure to acetylcholine reflects a difference in nicotinic receptor desensitization. To test this hypothesis, we treated the brain slice with the drug nicotine, which is well known to desensitize nicotinic receptors in the cortex (Quick and Lester, 2002; Paradiso and Steinbach, 2003; Picciotto et al., 2008), for 10 min at a concentration known to predominantly exert desensitizing effects in this neuronal population (100 nM; Bailey et al., 2010). The WT optogenetic cholinergic response was unchanged by application of nicotine; whereas the α5KO optogenetic response, was rapidly attenuated (Fig 4D-G). The cholinergic current charge measured in the WT and α5KO before and after the application of nicotine shows a significant interaction between the effect of nicotine and the genotype (2 way repeated measures ANOVA: Nicotine x genotype interaction: *F*_(1,16)_ = 9.8, \*\**p* = 0.006). Post hoc comparisons show that the WT response is not significantly different before and after nicotine (12 ± 1 pC vs 11 ± 2 pC, Sidak’s post hoc test: *t*_(16)_ = 0.61, *p* = 0.8; *N* = 7 mice) whereas the α5KO response is greatly reduced post nicotine (9 ± 0.5 pC vs 2 ± 0.6 pC, *t*_(16)_ = 5.39, \*\*\**p* < 0.001; *N* = 5 mice; Fig 4F). We conclude that the elimination of endogenous cholinergic responses in the α5KO following acute exposure to nicotine is due to desensitization of nicotinic receptors lacking the α5 subunit. Thus, *Chrna5* is essential to protect prefrontal endogenous cholinergic signaling from desensitization induced either by high acetylcholine levels or acute exposure to nicotine.

### Replication in a different optogenetic model: Chrna5 limits desensitization

We tested the robustness of our observation that α5KO endogenous cholinergic responses are more vulnerable to desensitization by examining whether it was independent of transgenic strategy for optogenetic release. Accordingly, we repeated experiments to measure nicotinic receptor desensitization in ChAT-IRES-Cre/+ Ai32/+ α5WT and α5KO mice, isolating the nicotinic response with atropine, and using the same accommodating-frequency optogenetic stimulus train used above. In these ChAT-IRES-Cre/+ Ai32/+ mice, the train response is stronger than in the ChATChR2 mice but is replicable and shows a significant genotype difference in the timing of its peak (*t*_(29)_ = 2.20, \**p* = 0.03), with α5KO peak occurring 37.2 ± 16.9 ms after the α5WT (*N* = 4 mice per genotype). The delay is observed in the absence of a difference in the peak cholinergic current (α5WT: 54 ± 6 pA vs α5KO: 51 ± 4 pA, unpaired t-test: *t*_(29)_ = 0.32, *p* = 0.7) or in the cholinergic charge transfer (α5WT: 21 ± 3 pC vs α5KO: 24 ± 2 pC, unpaired t-test: *t*_(29)_ = 0.90, *p* = 0.4). In short, the properties of the train responses in the ChAT-IRES-Cre/+ Ai32/+ α5WT and α5KO mice are suitable for testing whether loss of α5 *increases* vulnerability to nicotine desensitization.

For this experiment, the cholinergic response magnitude in voltage-clamp to a train of optogenetic stimulation was measured at different time points during the application of 100 nM nicotine (**Figure 5**). In this optogenetic line, there is again a significant interaction between the effect of nicotine exposure and genotype (2 way repeated measures ANOVA: nicotine x genotype interaction: *F*_(4,40)_ = 10.19, \*\*\*\**p* < 0.0001). Similar to results obtained in ChAT-ChR2 α5KO mice, we see that ChAT-IRES-Cre/+ Ai32/+ α5KO mice show almost-complete desensitization at the end of 10-minute exposure to nicotine (fraction of response at 5 min: 0.36 ± 0.06, Sidak’s post hoc test: *t*_(5)_ = 10.80, \*\*\**p* = 0.0005; at 10 min: 0.16 ± 0.04, *t*_(5)_ = 20.06, \*\*\*\**p* < 0.0001). While considerably milder, there is also significant nicotine-elicited desensitization in ChAT-IRES-Cre/+ Ai32/+ α5WT mice (fraction of response at 5 min: 0.61 ± 0.09, Sidak’s post hoc test: *t*_(5)_ = 4.20, \**p* = 0.03; at 10 min: 0.57 ± 0.08, Sidak’s post hoc test: *t*_(5)_ = 5.77, \*\**p* = 0.009). The stronger optogenetic release of acetylcholine in this optogenetic line helps to illustrate the degree to which *Chrna5* enables the α5WT to resist desensitization.

**Figure 5:**
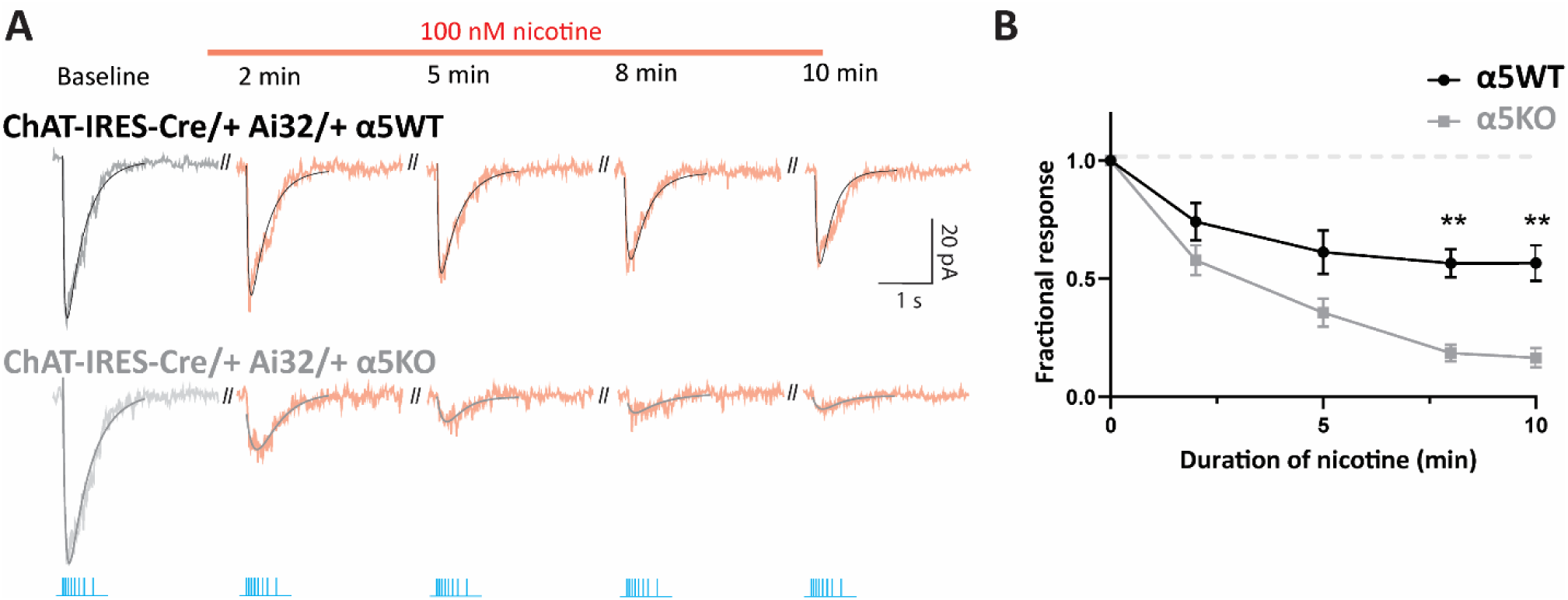
*Chrna5* attenuates desensitization of endogenous cholinergic signals in a different optogenetic model. **A**, Optogenetically evoked nicotinic responses in an α5WT and α5KO ChAT-IRES-Cre/+ Ai32/+ neuron at different time points during the application of 100 nM nicotine for 10 minutes. **B**, Time course of change in endogenous nicotinic response as nicotine is applied (2-way RM ANOVA: Genotype x nicotine interaction: *F*_(4,12)_ = 11.8, \*\**p* < 0.001; Sidak’s posthoc test comparing WT and α5KO: \*\**p* < 0.01).

These results confirm the critical role of *Chrna5* in protecting endogenous cholinergic signaling from desensitization. Together with the results in figures 1-4, we are able to show using two different transgenic strategies for optogenetic acetylcholine release that *Chrna5* in layer 6 of the prefrontal cortex has 2 roles: i) *Chrna5* is essential for a rapid onset of cholinergic activation and ii) *Chrna5* protects prefrontal endogenous cholinergic signaling from desensitization induced either by high acetylcholine levels or by acute exposure to nicotine.

### Rescuing rapid cholinergic response onset by targeting an unorthodox binding site

The α5KO mice show a deficit in the timing of cholinergic excitation that would delay input integration and the activation of postsynaptic partners, a finding consistent with the attention deficits observed in α5KO mice (Bailey et al., 2010). The manipulations that we have tested, including blocking presynaptic muscarinic autoinhibition (Fig 2) and inhibiting acetylcholinesterase (Fig 4), fail to rescue this slow onset of cholinergic excitation. Our optogenetic results suggest that attempts to rescue endogenous cholinergic signaling in α5KO mice must navigate a careful path between hastening response onset and avoiding desensitization. Therefore, we chose to examine a strategy of positive allosteric modulation, aiming to potentiate the nicotinic response without altering the duration of nicotinic receptor stimulation. Accordingly, we took advantage of the recently-identified α-α nicotinic receptor binding site and its selective agonist NS9283 (Grupe et al., 2013; Olsen et al., 2013, 2014; Jain et al., 2016). Stimulation of this unorthodox nicotinic receptor binding site does not produce a current in itself but instead enhances the affinity for acetylcholine at the orthodox α4-β2 binding sites (Wang and Lindstrom, 2018). Such enhancement in wildtype mice improves cognitive performance on attention tasks (Timmermann et al., 2012; Mohler et al., 2014).

Here, we asked if the unorthodox site agonist NS9283 can rescue the onset kinetics of the endogenous cholinergic response in α5KO mice without triggering desensitization. One requirement for this approach to work is that at least a proportion of nicotinic receptors in α5KO must have adopted the (α4)_3_(β2)_2_ stoichiometry (**Figure 6A**). We measured cholinergic responses to a train of optogenetic stimulation in ChAT-ChR2 α5KO mice in the presence of atropine before and after the application of 100 nM NS9283 for 5 minutes (Fig 6B-E). We found that this minimal concentration of NS9283 is highly effective at speeding the onset of cholinergic activation (increase in rising slope: 80 ± 30 pA/s, *N* = 3 mice, paired t-test: *t*_(8)_ = 2.66, \**p* = 0.03; Fig 6C). NS9283 also increased the peak amplitude of the response (change: 12 ± 2 pA, paired t-test: *t*_(9)_ = 4.99, \*\*\**p* = 0.0008). Further assessment of the restorative capacity of NS9283 in current-clamp again showed that it significantly increased the onset speed of cholinergic responses (increase in rising slope: 12 ± 5 mV/s, paired t-test: *t*_(9)_ = 2.41, \**p* = 0.04; Fig 6E) and the response amplitude (change: 4 ± 1 mV, paired t-test: *t*_(9)_ = 5.18, \*\*\**p* = 0.0006). Figure 6F shows example exponential fits to cholinergic responses in a WT, α5KO and the same α5KO neuron after application of 100 nM NS9283. We see that the application of NS9283 is able to rescue the slow onset kinetics of α5KO cholinergic responses to match the WT response. Furthermore, we observe that the rescue of onset kinetics and potentiation caused by NS9283 is long-lasting without triggering significant desensitization of nicotinic receptors upon subsequent cholinergic stimulation (Fig 6G). There is no significant effect of continuous NS9283 application on the amplitude of cholinergic responses obtained in α5KO (one-way repeated measures ANOVA: F_(1.9, 3.9)_ = 2.74, *p* = 0.18). Thus, using a low concentration of NS9283, we are able to rescue the onset of cholinergic responses in α5KO to achieve the rapid timing observed in WT without engaging desensitization mechanisms.

**Figure 6:**
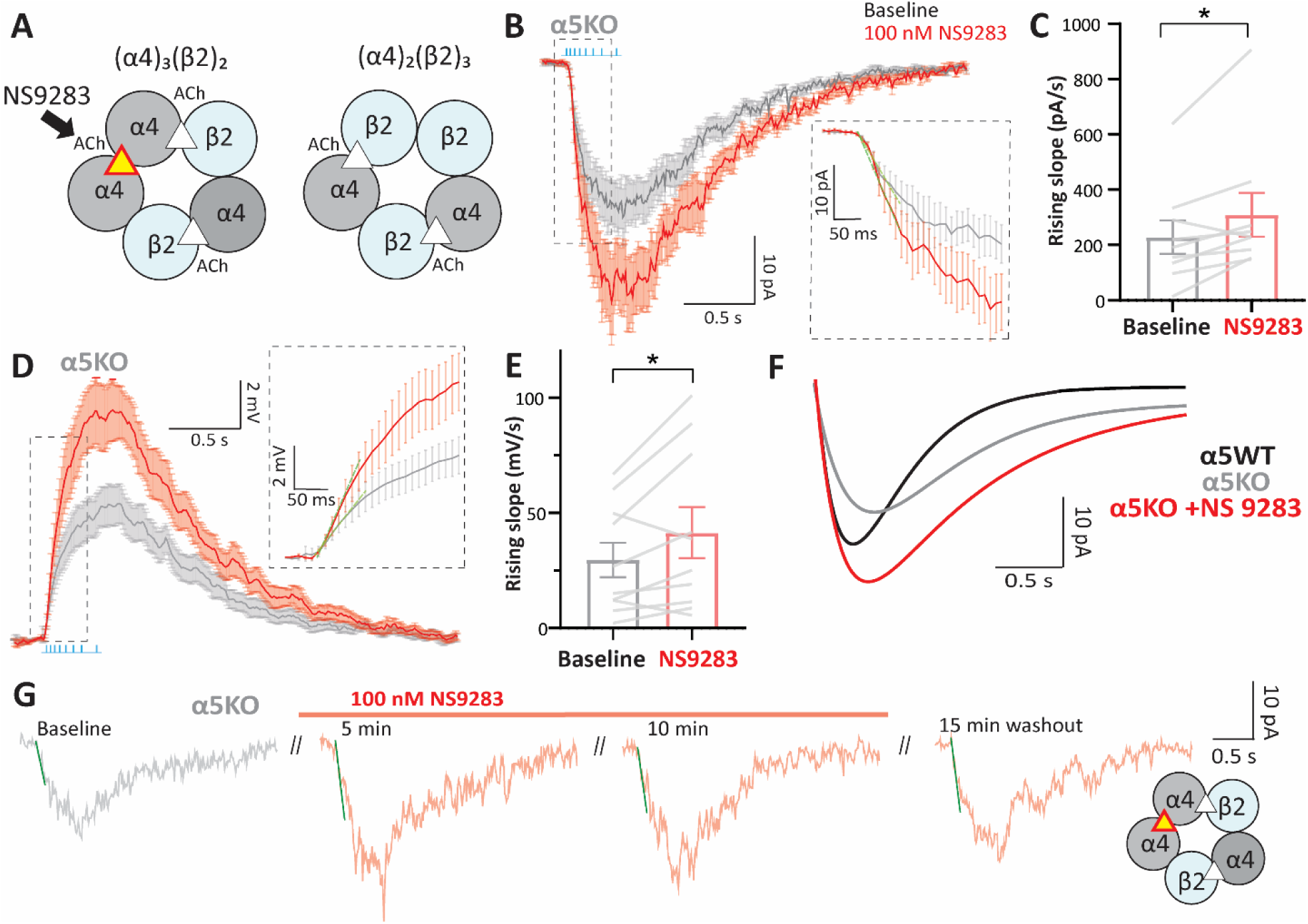
Pharmacological restoration of rapid kinetics of endogenous cholinergic responses after disruption of *Chrna5*. **A**, Schematic illustrating the two possible stoichiometries of nicotinic receptors in the α5KO. NS9283 is a selective agonist for the α-α site found in (α4)_3_(β2)_2_ receptors. **B, D**, Average current (B) and depolarization (D) in response to optogenetic acetylcholine release before and after the application of 100 nM NS9283 in the α5KO (*n* = 10 cells). Atropine was present throughout to isolate the nicotinic response. Inset: Fast rising phase of the response with linear fit (green) to the first 50ms of the response from light onset **C, E**, Bar graph showing the rising slope before and after NS9283 for (C) the current (paired t-test: *t*_(8)_ = 2.66, \**p* = 0.03) and (E) the depolarization (paired t-test: *t*_(9)_ = 2.41, \**p* = 0.04) determined from the linear fit in α5KO. NS9283 causes a significant increase in onset speed of cholinergic responses in α5KO. **F**, Example exponential fits to cholinergic responses in a WT, α5KO and the same α5KO neuron after application of 100nM NS9283. Application of NS9283 rescues slow onset kinetics of α5KO cholinergic responses to match the WT. **G**, Optogenetically-evoked nicotinic responses along with linear fits of the same α5KO neuron shown in (F) at 5 and 10 minutes of NS9283 application and following a 15-minute washout period. Potentiation caused by NS9283 is long-lasting and optogenetic release of endogenous acetylcholine can be repeated without triggering desensitization of nicotinic receptors.

## Discussion

Our results reveal that the α5 subunit encoded by *Chrna5* is necessary to generate rapid onset of responses to endogenous acetylcholine released optogenetically. In this way, it regulates the timing of the peak cholinergic modulation of layer 6 pyramidal neurons in prefrontal cortex, but not its magnitude. In addition, the α5 subunit protects the endogenous cholinergic signaling from desensitization induced by prolonged exposure to acetylcholine or acute nicotine. Finally, we show that the slow onset of cholinergic responses in mice lacking the α5 subunit can be rescued using NS9283, a selective agonist for the unorthodox α-α binding site on (α4)_3_(β2)_2_ nicotinic receptors.

### Chrna5 permits a rapid response to endogenous cholinergic signaling

Rapid cholinergic modulation of the prefrontal cortex is critical for attention. Layer 6 pyramidal neurons are key players in this phenomenon, since a large proportion are corticothalamic and can exert a direct top-down influence on incoming sensory inputs (Kassam et al., 2008; Thomson, 2010). Layer 6 corticothalamic neurons express the α5 nicotinic receptor subunit encoded by *Chrna5* which is critical for the performance of demanding attention tasks (Bailey et al., 2010). We observe that neurons lacking α5 showed significantly impaired kinetics in responding to endogenous acetylcholine release, exhibiting a much slower rise and delayed time of peak. The slow onset of cholinergic activation in α5KO results in a delay of up to ∼100 ms in initiation of acetylcholine induced spiking in layer 6 neurons. We posit that the delay in cholinergic activation in the α5KO could result in failure to integrate inputs or activate postsynaptic targets within a critical window critical for the detection of sensory cues, leading to attention deficits observed in these mice (Bailey et al., 2010).

### Temporal constraints on endogenous cholinergic signaling

We demonstrate that the cholinergic inputs to the layer 6 pyramidal neurons are under strong M2 muscarinic receptor mediated autoinhibition of release (∼ 40% suppression of the full response by active presynaptic M2 receptors) in both WT and α5KO. Releasing the autoinhibitory brake on acetylcholine release does not improve the aberrant cholinergic kinetics in the α5KO. However, the strong muscarinic autoinhibition of endogenous cholinergic release onto layer 6 pyramidal neurons would be predicted to restrict the emphasis onto the fast-rising phase of a response to a train of cholinergic stimuli. The acetylcholine released to the first stimulus will activate the presynaptic M2 receptors and suppress further release due to the subsequent stimuli. The potential for such autoinhibition highlights the importance of the α5 nicotinic subunit in generating an initial rapid response to the acetylcholine release. Together with the high expression of the metabolic enzyme acetylcholinesterase in deep layers of the prefrontal cortex (Sendemir et al., 1996; Anderson et al., 2009), our results suggest that prefrontal layer 6 cholinergic modulation is hardwired for rapid transient effects with *Chrna5* ensuring rapid postsynaptic activation.

### Cognitive ramifications of rapid cholinergic signaling

Acetylcholine release in the cortex has been shown to vary on rapid timescales with behavioural state-cholinergic axon activation in the barrel cortex correlates rapidly with whisking behaviour (Eggermann et al., 2014). Similarly, activity of cholinergic axons in the auditory cortex rapidly shifts and is predictive of behavioural context (Kuchibhotla et al., 2017). Fast cholinergic transients are observed in the prefrontal cortex in association with rewards, and cue detection on sustained attention tasks (Parikh et al., 2007; Gritton et al., 2016; Teles-Grilo Ruivo et al., 2017b). Prefrontal nicotinic receptors are required for the initial transition from low gamma to high gamma states coinciding with cue presentation in an attention task (Howe et al., 2017). The attention deficits observed in mice lacking *Chrna5* performing a 5 choice serial reaction time test were also critically dependent on timing (Bailey et al., 2010): α5KO mice were impaired only at the briefest and most-demanding stimulus durations. The slower cholinergic activation of layer 6 corticothalamic neurons in α5KO would be consistent with a failure to detect brief cues within a critical window for integration.

### Chrna5 to protect the synaptic cholinergic response

While rapid cholinergic signaling in the PFC is critical for detection of sensory cues, cholinergic tone in the PFC is important under challenging conditions of sustained attention (Sarter and Lustig, 2019, 2020). High cholinergic tone in the PFC is associated with sustained attention and top down attentional control in the presence of distractor challenges, and can well last beyond the task duration (Himmelheber et al., 2000; St. Peters et al., 2011; Paolone et al., 2012). Prefrontal acetylcholine levels also greatly increase during conditions requiring high cognitive effort and stress (Mark et al., 1996; Pepeu and Giovannini, 2004; Teles-Grilo Ruivo et al., 2017a). To replicate this scenario *ex vivo*, we prolonged acetylcholine presence by blocking acetylcholinesterase irreversibly and examined the role of *Chrna5*. This experiment revealed a sharp dichotomy between the genotypes, where cholinergic responses after acetylcholinesterase block were much smaller in the α5KO compared to the WT. Furthermore, acute exposure to a low level of nicotine thought to mimic the concentrations seen in smokers (Rose et al., 2010) sharply attenuated synaptic cholinergic transmission in the α5KO, while WT cholinergic transmission was resilient, revealing that the α5 nicotinic subunit has a critical role in protecting against desensitization. These experiments demonstrate using endogenous acetylcholine release to physiological stimulation patterns, a critical role for the α5 subunit in conferring a protective role against desensitization during elevated cholinergic tone or acute nicotine exposure.

### A novel treatment approach and clinical relevance

The loss of *Chrna5* causes profound attention deficits (Bailey et al., 2010; Howe et al., 2018) and it is of great interest to identify pharmacological interventions to correct this dysfunction. However, the vulnerability of α5KO animals to complete desensitization of their endogenous cholinergic signaling is of utmost importance when considering approaches to treat the attention deficits with cholinergic modulators. Treatment with cholinesterase inhibitors in animals lacking the α5 subunit is problematic as it could engage powerful desensitization of endogenous prefrontal cholinergic signaling. We instead show that aberrant cholinergic kinetics which may underlie attention deficits in α5KO animals can be rescued partly by NS9283, an agonist for the unorthodox α-α binding site, that allosterically enhances nicotinic receptor affinity without causing desensitization. A low concentration of NS9283 was able to restore the slow onset of synaptic cholinergic responses in α5KO to WT levels. NS9283 has been previously shown to improve attentional performance in wildtype animals, pointing to an underestimated potential of this drug to improve attention in compromised states (Timmermann et al., 2012; Mohler et al., 2014). Our work provides a novel pharmacological target-NS9283 which could be used to physiologically manipulate endogenous cholinergic signaling to improve attention in pathological states. This may be particularly relevant for the treatment of attention disorders in humans carrying prevalent non-functional polymorphisms in the *Chrna5* gene (Bierut et al., 2008).

Recent examinations of the cholinergic system have shown great interest in mechanisms underlying diverse spatiotemporal scales of cholinergic signaling in the cortex (Disney and Higley, 2020; Sarter and Lustig, 2020). Our study reveals a specialized role of the α5 nicotinic receptor subunit in generating the rapid cholinergic modulation of the prefrontal cortex known to be critical for cognition. Such kinetic properties may define critical windows for cognitive processing. We also show that the α5 nicotinic subunit protects rapid cholinergic signaling from desensitization induced by elevated acetylcholine levels or nicotine exposure. Finally, we demonstrate that rapid cholinergic signaling can be rescued in the absence of α5 without triggering desensitization by allosterically enhancing nicotinic receptors with NS9283, an agonist for the unorthodox binding site. Together, this work improves our understanding of cholinergic modulation of attention circuits and identifies a pharmacological target to restore the rapid kinetics of cholinergic signaling in pathological conditions.

## Acknowledgements

This research was funded by the Canadian Institutes of Health Research (CIHR MOP 89825, EKL), the Canada Research Chair in Developmental Cortical Physiology (EKL), and the Mary Beatty Fellowship from the University of Toronto (SV). We thank Ms. Janice McNabb and Mr. Ha-Seul Jeoung for expert technical assistance. Earlier presentations of this work received valuable feedback from Dr. Junchul Kim, Dr. Beverly Orser, and Dr. Steve Prescott of the University of Toronto.

